# Advanced image generation for cancer using diffusion models

**DOI:** 10.1101/2023.08.18.553859

**Authors:** Benjamin L. Kidder

**Author notes:** Correspondence: Benjamin L. Kidder.

## Abstract

Deep neural networks have significantly advanced medical image analysis, yet their full potential is often limited by the relatively small dataset sizes. Generative modeling has stimulated attention for its potential applications in the synthesis of medical images. Recent advancements in diffusion models have exhibited a remarkable capacity for producing photorealistic images. Despite this promising development, the application of such models in the generation of medical images remains underexplored. In this study, we explored the potential of using diffusion models to generate medical images, with a particular emphasis on producing brain magnetic resonance imaging (MRI) scans, such as those depicting low-grade gliomas. Additionally, we examined the generation of contrast enhanced spectral mammography (CESM) images, as well as chest and lung X-ray images. Utilizing the Dreambooth platform, we trained stable diffusion models based on text prompts, class and instance images, subsequently prompting the trained models to produce medical images. The generation of medical imaging data presents a viable approach for preserving the anonymity of medical images, effectively reducing the likelihood of patient re-identification during the exchange of data for research. The findings of this study reveal that the application of diffusion models in generating images successfully captures attributes specific to oncology within imaging modalities. Consequently, this research establishes a framework that harnesses the power of artificial intelligence for the generation of cancer medical imagery.

## INTRODUCTION

Generative diffusion models have generated considerable interest due to their facilitation of cutting-edge image generation techniques. These models have been developed utilizing extensive, multi-modal data sets, such as LAION-5B^1^, which include images and their corresponding textual descriptions. There has also been a surge in interest surrounding generative models in the field of medical imaging^2–14^. The primary aim of these methodologies is to generate data that emulates the ground truth imaging data with clinical features. This generated data holds the potential to enhance the efficacy of a myriad of subsequent processes, thereby eliminating or reducing the need for acquiring extensive, expensive expert-annotated data sets for rare clinical cases.

Generative Adversarial Networks (GANs) have paved the way for innovative strategies to tackle a wide range of complex pathological image analysis tasks^15–27^. Despite their potential, GAN models are susceptible to overfitting, ultimately leading to suboptimal image synthesis outcomes. For image generation, leading-edge models like DALLE-2^28^, Mid-Journey, and Stable Diffusion^29^ have gained significant recognition. Of these, Stable Diffusion is unique in providing open-source code. Stable diffusion follows the pioneering work of text-to-image diffusion models GLIDE^30^, Imagen^31^, and DALLE-2^28^. The growing popularity of diffusion models^28–31^, also known as diffusion probabilistic models^32^, in contrast to GANs^33–36^, is largely due to their superior stability throughout the training phase.

Leveraging image generation methodologies presents a promising avenue for addressing the paucity of meticulously curated, high-resolution medical imaging datasets^37^. The annotation process for such datasets requires substantial manual labor from skilled medical professionals, who possess sufficient expertise to decipher and attribute semantically significant features within the images. This innovative approach alleviates the burden on healthcare professionals while ensuring the availability of reliable data.

Here, we utilized Dreambooth^14^, which is dependent on the Imagen^31^ text-to-image model, by employing a version that has been integrated with the Stable Diffusion framework. Providing a minimal number of medical images as input, we refine a pre-trained text-to-image stable diffusion model to associate a distinct identifier with the specific medical imaging modality. Through the association of a distinct identifier with medical images and embedding them within the model, this method enables the generation of new, photorealistic medical images for cancer and other diseases. These high-quality synthetic datasets hold the potential to supplement traditional data acquisition methods, particularly in cases where medical imaging data is scarce or inadequate. These medical imaging datasets hold immense potential to revolutionize the landscape of medicine, by effectively addressing data scarcity and driving significant advancements in disease detection and treatment. Furthermore, these datasets can serve as resources for machine learning strategies.

## RESULTS

### Training of the Dreambooth stable diffusion framework for brain cancer

The experiments were performed using Dreambooth^14^ integrated with the stable diffusion^29^ v1.5 framework. The process involves gradually adding Gaussian noise to an image and then denoising it to generate new images (**Fig. 1**). Nvidia V100 or A100 GPUs were utilized to perform stable diffusion. Model weights (runwayml/stable-diffusion-v1-5) were obtained from the Huggingface platform^38^, which utilizes the diffusers library^39^. The settings for training an AI model include a pretrained model and VAE (stabilityai/sd-vae-ft-mse). It employs prior preservation with a weight of 1.0 and sets a specific seed value (1337) and resolution (512). With a training batch size of 1, the text encoder is trained using mixed precision, an 8-bit Adam optimizer, and one gradient accumulation step. The learning rate is set to 1e-6, and a constant scheduler is used with no warmup steps. The model utilizes 50 class images, a sample batch size of 4, a maximum of 800 training steps, and a save interval of 10,000.

**Figure 1.**
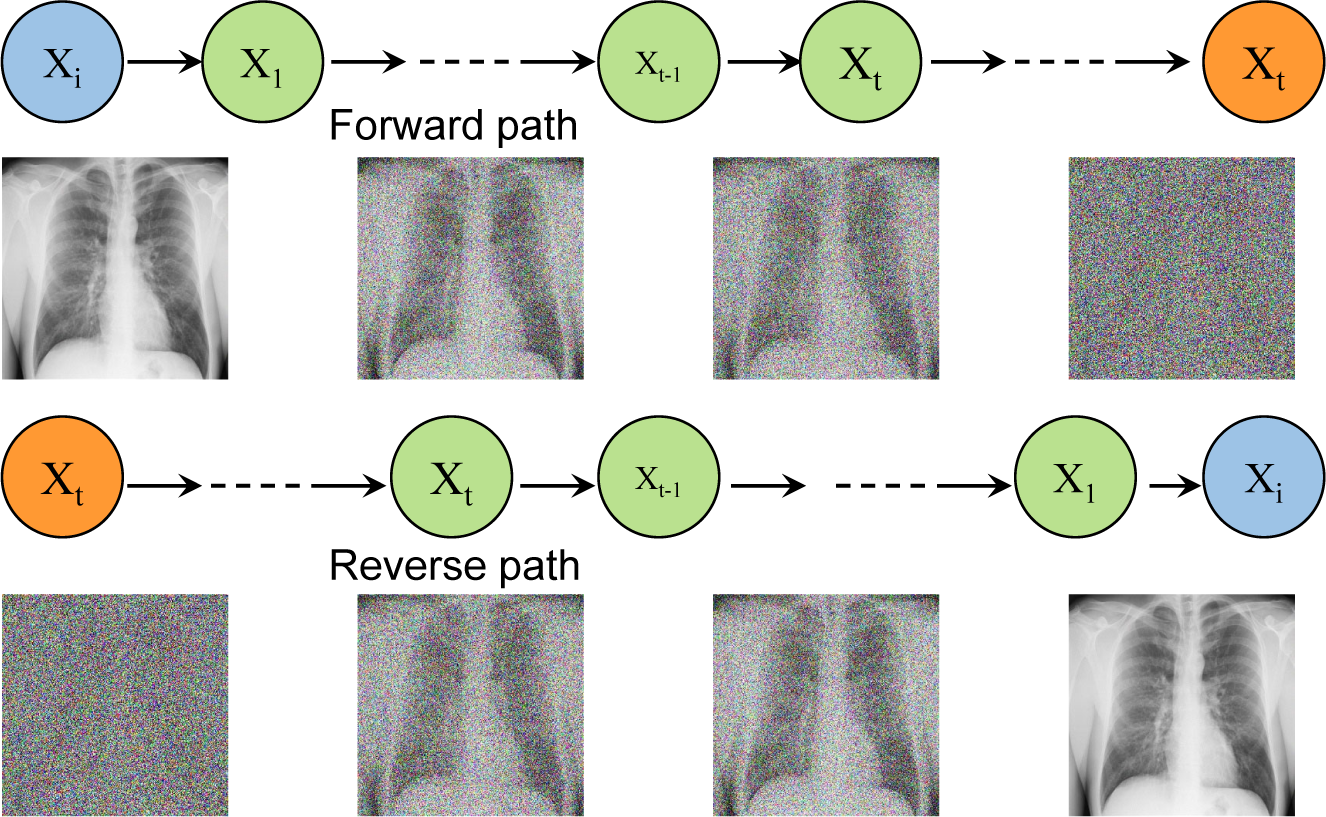
Schematic of the diffusion model training process. Schematic of the dreambooth stable diffusion process featuring forward and reverse diffusion^5,57^.

Using this approach, we trained stable diffusion models using clinically curated medical images for various types of brain cancer. This includes tumor classifications featured in the Kaggle dataset, “Brain Tumor Image Dataset,” which includes images from gliomas, meningiomas, and pituitary tumors^40,41^. We then synthesized images using the trained model by inputting textual prompts. The collection of representative synthesized brain cancer images features MRI scans illustrating meningiomas (**Fig. 2A**), gliomas (**Fig. 2B**), and pituitary tumors (**Fig. 3**). Additionally, we have generated images showcasing sagittal and cross-sectional MRI brain scans following training of images from healthy brains (**Fig. 4**).

**Figure 2.**
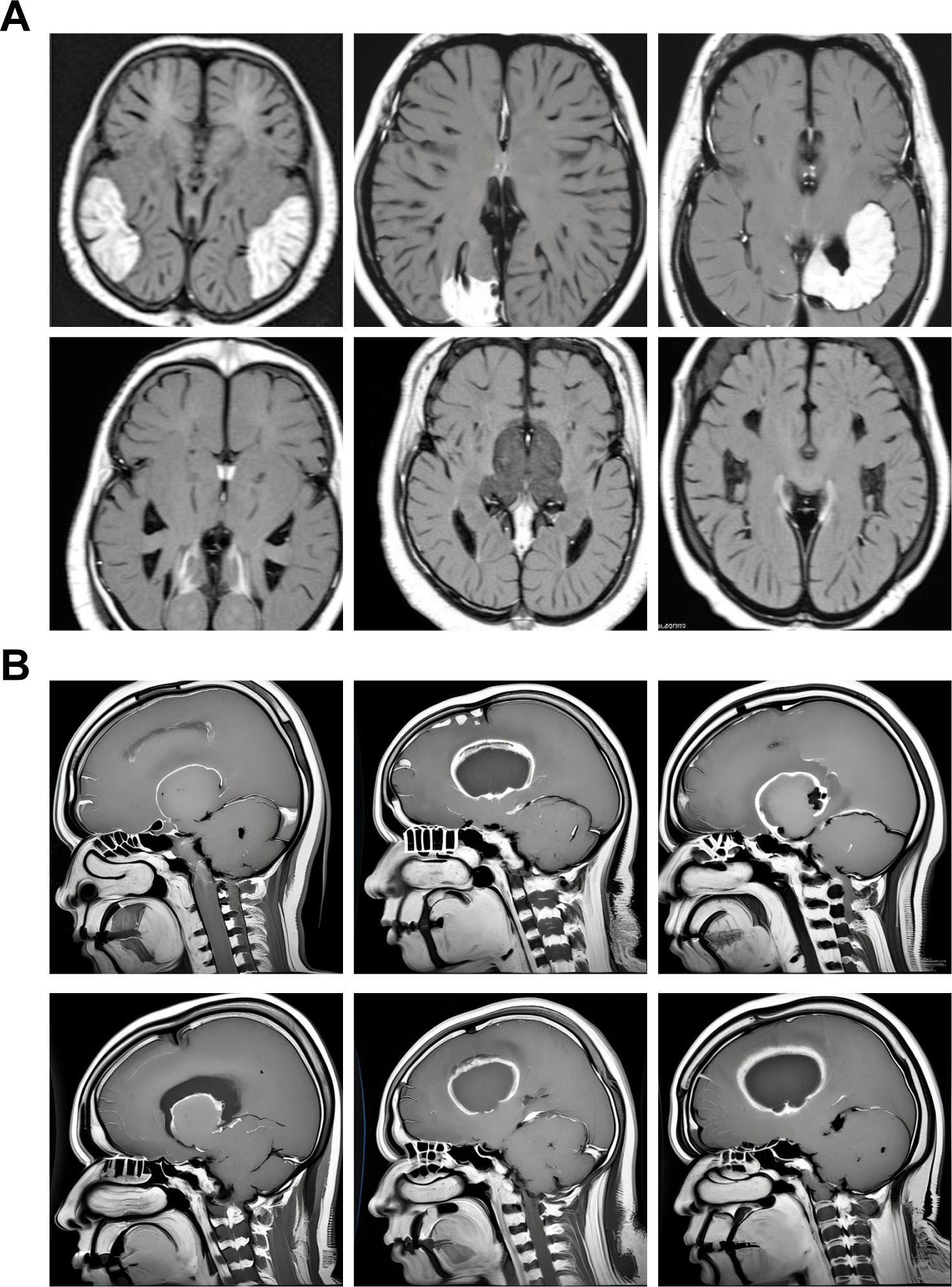
Visualization of MRI images of brain cancer generated by fine-tuning a Dreambooth diffusion model. (A) Cross-sectional MRI scans of synthesized meningioma tumors, displaying z-slices of the brain. (B) Sagittal brain MRI scans of glioma tumors generated using a stable diffusion model fine-tuned with the Kaggle “Brain Tumor Image Dataset” containing gliomas, meningiomas, and pituitary tumors. Images have a resolution of 512×512 pixels.

**Figure 3.**
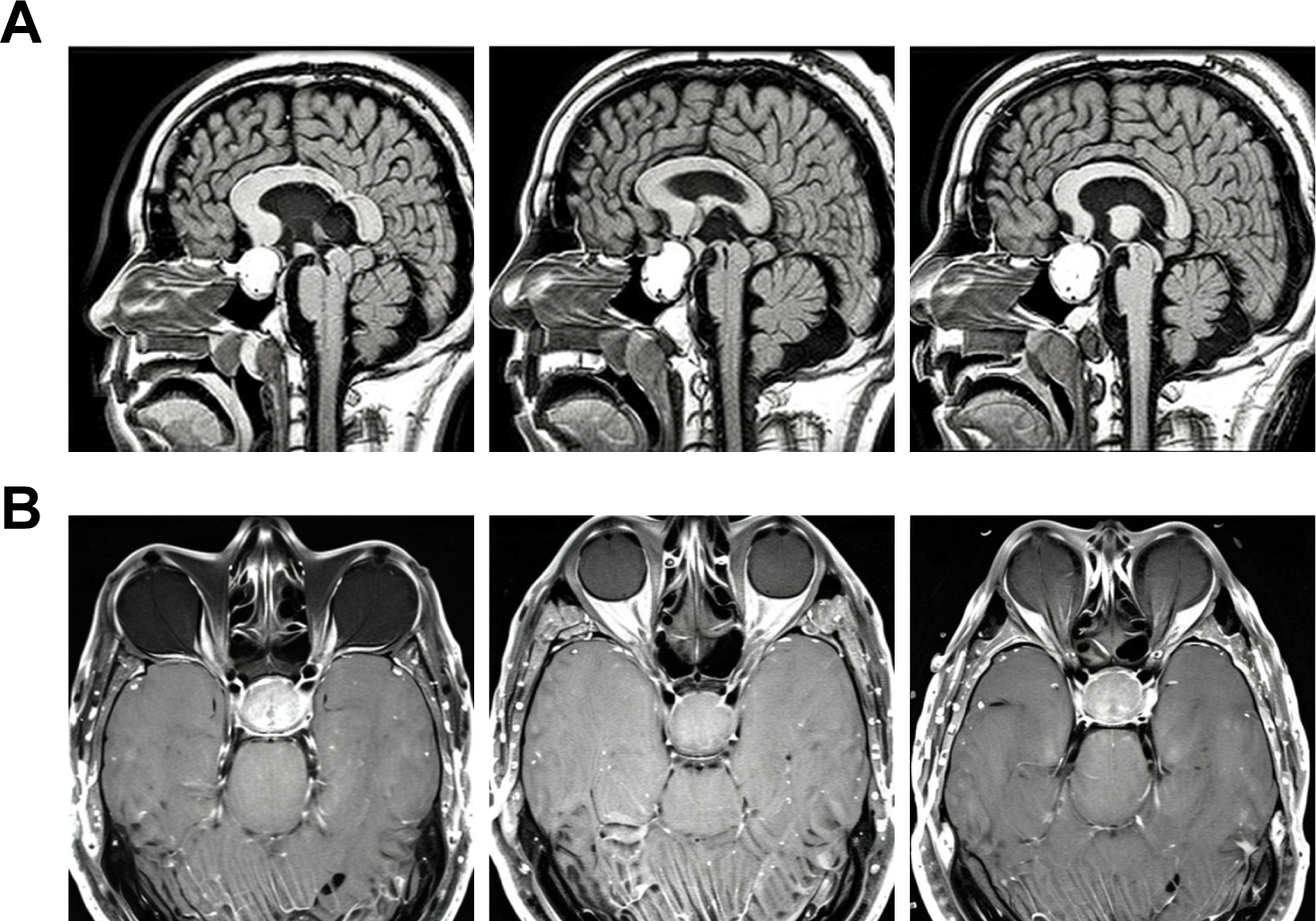
MRI images depicting pituitary tumor brain scans produced through refining a Dreambooth diffusion model. (A) Sagittal sectional MRI scans displaying pituitary tumors and brain tissue, with each micrograph depicting a slice of the head, brain, and tumor. (B) Cross-sectional brain MRI scans of pituitary tumors produced using a stable diffusion model refined with the Kaggle “Brain Tumor Image Dataset,” which includes gliomas, meningiomas, and pituitary tumors. Images are displayed at a resolution of 512×512 pixels.

**Figure 4.**
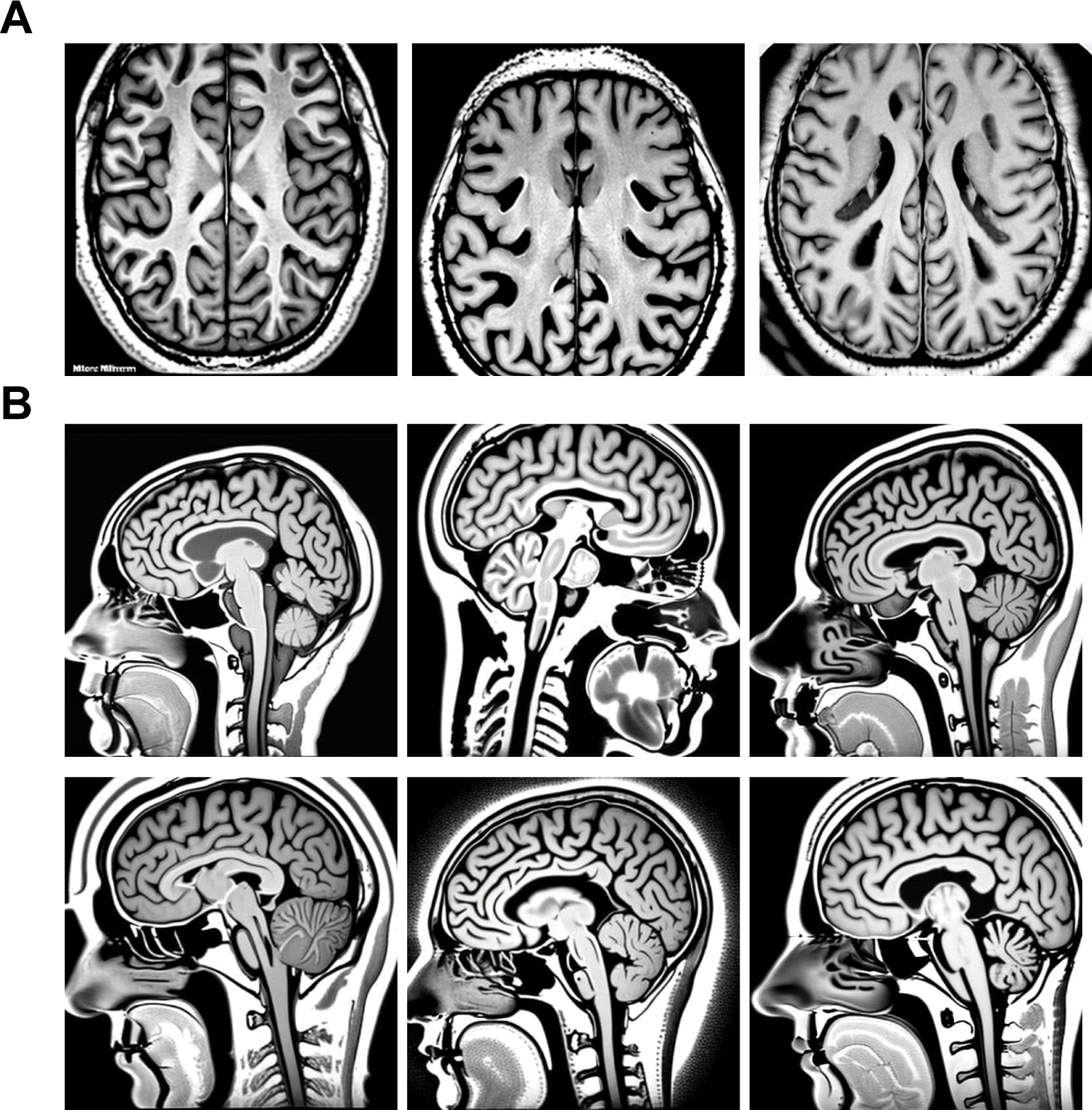
Examples of images produced for healthy brain MRI scans. (A) Synthesized images of brain MRI scans in cross-sectional view, showing pituitary tumors and brain tissue, where each micrograph represents a section of the head, brain, and tumor. (B) Synthesize MRI scans of a healthy brain in sagittal sections. These images were produced using a dreambooth stable diffusion model and feature a resolution of 512×512 pixels.

Low-grade glioma (LGGs), such as those classified by the WHO as grades II and III, are a collection of brain neoplasms^42–44^. Unlike grade I gliomas, which can frequently be cured through surgical removal, grades II and III gliomas display infiltrative characteristics, making them prone to recurrence and progression into more advanced forms^45^. Relying on histopathological information to predict the prognosis for individuals with these tumor types is imprecise and subject to variation among different observers. A potential solution to this challenge could be the classification of LGG subtypes by means of image examination and clinical evaluation. Imaging techniques offer valuable insights prior to surgical intervention or when tumor removal is unattainable. Studies have unveiled a connection between the morphological characteristics of tumors derived from MRI data and their genomic classifications^46^. However, the initial process of obtaining these tumor features necessitated labor-intensive, expensive manual segmentation of the MRI scans^47^. This method leads to manual annotations that exhibit variability.

As a result, we have developed LGG diffusion models to aid the scientific community in refining algorithms for the categorization of gliomas and various other malignancies. To this end, we have generated stable diffusion models of lower-grade gliomas using Dreambooth and medical images from the LGG Segmentation Dataset^48^, which reveal diversity of MRI images following text-to-image prompting (**Fig. 5**).

**Figure 5.**
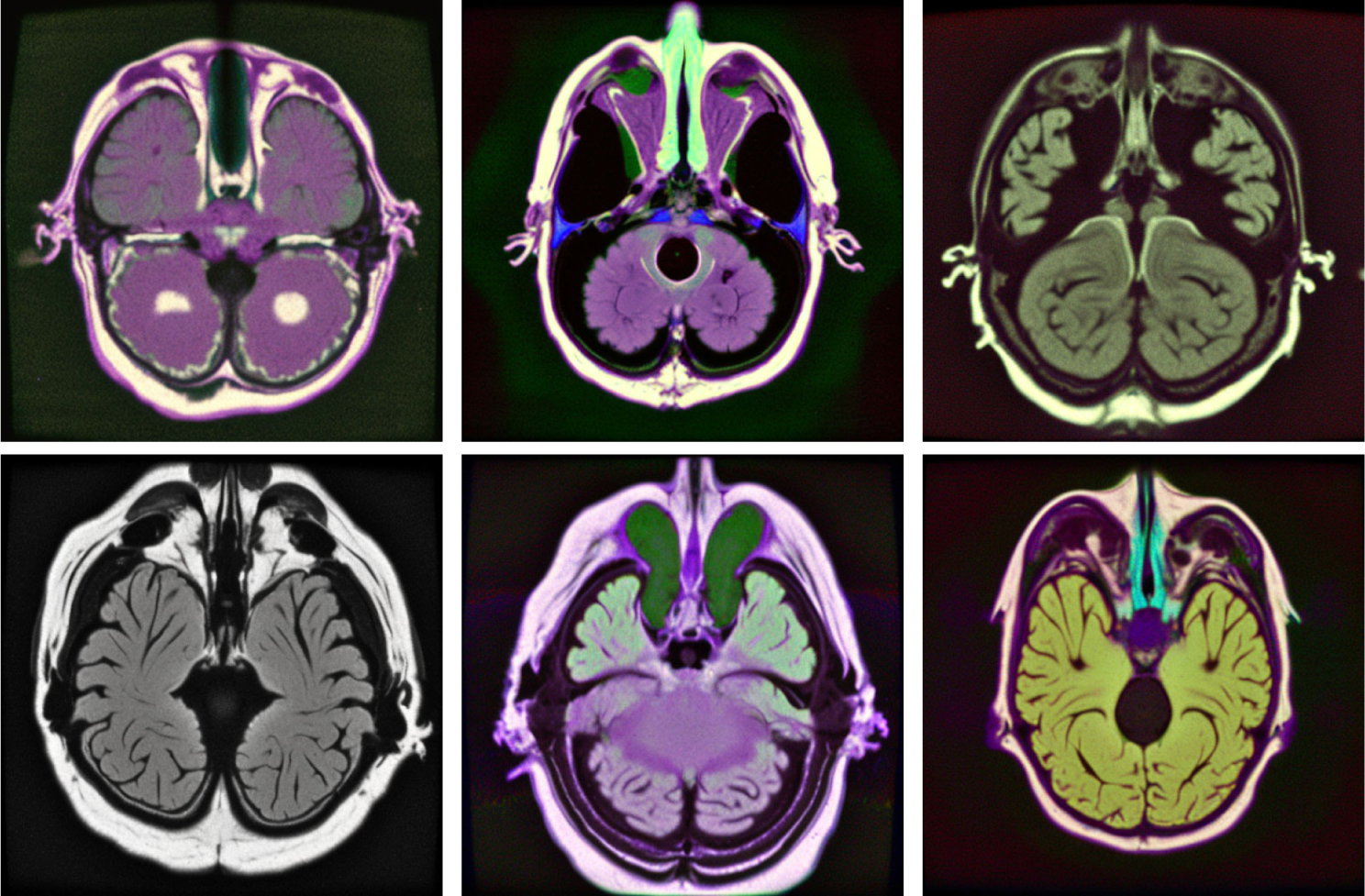
Images generated for lower-grade gliomas. Images utilized for fine-tuning a dreambooth stable diffusion model were sourced from the Cancer Genome Atlas Low Grade Glioma Collection (TCGA-LGG) and the LGG segmentation datasets available on Kaggle (https://www.kaggle.com/datasets/mateuszbuda/lgg-mri-segmentation). These images were acquired from The Cancer Imaging Archive (TCIA) and are associated with patients in TCGA lower-grade glioma collection. The resulting synthesized images are displayed at a resolution of 512 x 512 pixels.

### Generative models for mammography images

The application of deep learning presents an opportunity to alleviate the burden on radiologists while simultaneously enhancing diagnostic accuracy. However, extensive, and comprehensively annotated datasets are essential to realize these goals. The categorized Digital Database for Low energy and Subtracted Contrast Enhanced Spectral Mammography images (CDD-CESM)^49^ dataset comprises 2,006 high-resolution annotated images from contrast-enhanced spectral mammography (CESM) accompanied by medical records from patients aged 18-90. Utilizing conventional digital mammography devices, CESM was conducted in the study with software that enabled image capture.

Using data from the CDD-CESM repository^49^, we trained a Dreambooth stable diffusion model using a minimal number of CESM mammography images. Subsequently, we utilized textual prompts to produce synthesized mammography images (**Fig. 6A**). Additionally, we utilized image-to-image diffusion to create a diverse array of breast cancer mammography images, thereby minimizing patient-specific traits and generating original breast tumor images (**Fig. 6B**). Fine-tuning of these stable diffusion models and image-to-image algorithms enable the generation of images exhibiting distinct clinical attributes, making them suitable for various purposes such as medical education, machine learning, and the procurement of images related to rare conditions. Additionally, these images can be employed in diagnostic assistance, therapeutic planning, patient education, and the development of novel imaging technologies.

**Figure 6.**
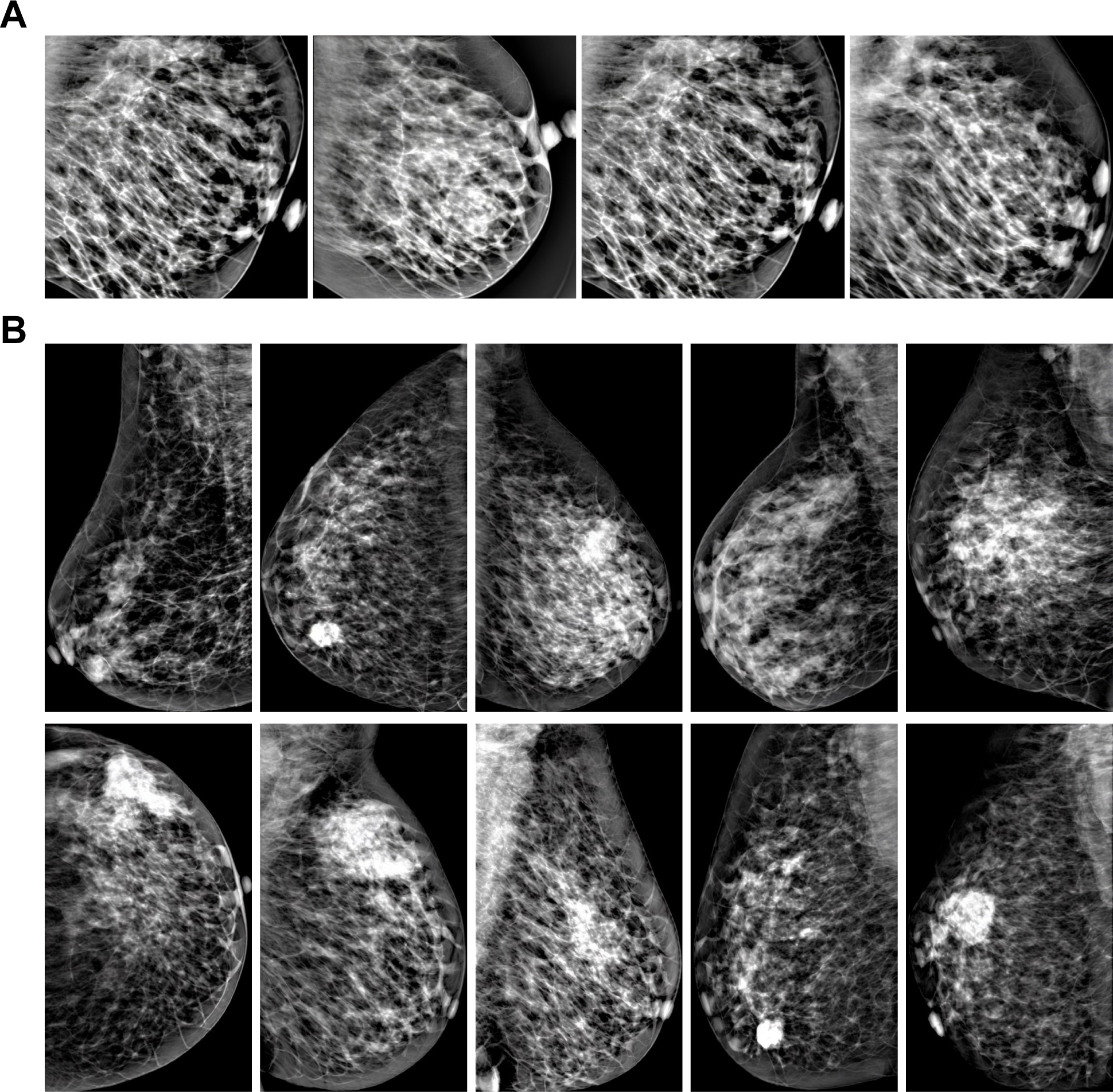
Synthesis of contrast-enhanced spectral mammography (CESM) images. (A) Training images were obtained from the Categorized Digital Database for Low Energy and Subtracted Contrast Enhanced Spectral Mammography (CDD-CESM)^49^ database to develop a robust diffusion model. The fine-tuned model produced synthetic mammography images. (B) Text-to-image synthesis of mammography images using stable diffusion enabled the generation of diverse mammography images for breast cancer detection. Images are presented at a resolution of 512×512 pixels.

### Synthesis of chest x-rays images using diffusion models

The thoracic radiograph is a prevalent diagnostic tool employed for the identification and assessment of numerous pulmonary disorders, including pneumonia^50^, lung cancer^51^, chronic obstructive pulmonary disease (COPD)^52^, interstitial lung diseases^53^, and pulmonary embolism^54,55^. X-ray images, along with their corresponding radiology evaluations, are generated and archived within the databases of healthcare facilities. We utilized the thoracic radiograph repository to develop a fine-tuned generative diffusion model to create synthesized chest X-rays using a few images. This model utilized images from the ChestX-ray8 database^56^, which encompasses 108,948 anterior perspective X-ray images from 32,717 distinct individuals. Subsequently, we employed textual prompts to generate synthesized thoracic radiographs (**Fig. 7**). The resulting images showcase detailed chest X-ray visuals, illustrating the application of the trained diffusion model in the creation of pulmonary and chest radiographic images. Synthesized chest images for lung and pulmonary diseases provide numerous advantages, including enhanced machine learning training data and preserved patient privacy. These cost-effective images may serve as valuable educational resources and allow researchers to create customized data for innovative diagnostic and treatment approaches. Additionally, using synthesized images ensures ethical research practices and unbiased evaluations of diagnostic algorithms.

**Figure 7.**
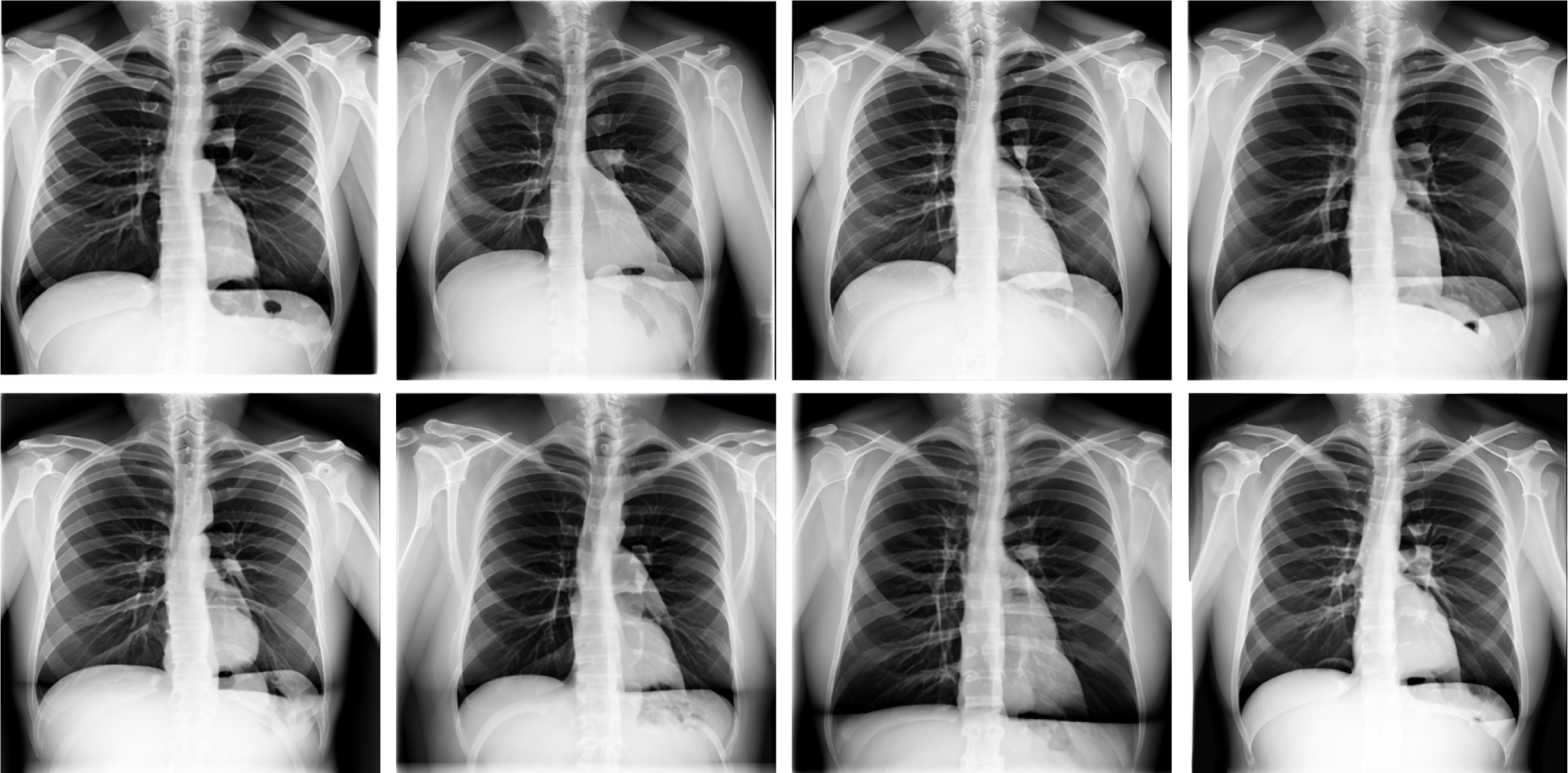
Chest x-ray images synthesized using a fine-tuned diffusion model. Synthesis of chest X-ray images through the application of Dreambooth Stable Diffusion for text-to-image transformation, which is employed for model fine-tuning. A retrained or fine-tuned Dreambooth model denoises from a random Gaussian noise vector, conditioned by embeddings generated from text prompts. The decoder within the models variational autoencoder maps the noise-reduced latent vector to pixel space, yielding high-quality, diverse chest X-rays.

## DISCUSSION

By utilizing Dreambooth stable diffusion and a diverse set of medical imagery, we have developed diffusion models. The use of stable diffusion to create medical images could lead to progress in areas such as diagnostics, research, and the development of treatments. The production of high-quality medical images may lead to enhanced diagnostic precision, enabling healthcare professionals to accurately identify and assess various medical conditions. Creating medical images can be beneficial for developing algorithms and machine learning models that enhance diagnostic capabilities and treatment strategies. Also, generating medical images promotes a more comprehensive understanding of human diseases and tumor biology, helping to unravel the complex mechanisms underlying cancer development and progression. This knowledge is crucial for identifying potential therapeutic targets and designing effective treatments.

Utilizing synthesized chest images for lung and pulmonary diseases offers a range of benefits. These images can enhance training data for machine learning algorithms, leading to improved diagnostic capabilities while preserving patient privacy. Moreover, generating synthetic images is cost-effective, as it reduces the need for expensive medical imaging equipment and reliance on limited real-world data. Synthesized images also serve as valuable educational tools, helping medical students and professionals better understand and identify various lung and pulmonary diseases. Customized datasets can be created with specific disease characteristics, promoting the development and testing of innovative diagnostic methods and treatment approaches. The use of synthesized images ensures ethical research practices by eliminating the need for invasive procedures on real patients, and carefully curated synthetic datasets can avoid biases, providing a more balanced and accurate evaluation of diagnostic algorithms and techniques.

The potential of Dreambooth stable diffusion models for mammography images has also been showcased in this study. This research may be beneficial for developing diagnostic tools and training medical professionals. The synthesized mammography images provide a rich dataset to study various breast cancer cases, including rare conditions that may not be easily accessible in clinical settings. Image synthesis offers a valuable resource for creating a diverse range of mammography images, enabling the development of more accurate diagnostic tools. By leveraging these synthesized images, machine learning algorithms can be trained and fine-tuned, ultimately contributing to improved diagnostic accuracy and a reduced burden on radiologists. Furthermore, these images can serve as a basis for the creation of innovative diagnostic techniques, broadening the scope of medical imaging and enhancing patient care. This information may contribute to a more comprehensive understanding of breast cancer diagnosis and treatment, ultimately resulting in improved patient outcomes.

Despite the promising findings of this study, future research should address limitations such as the generalizability of the results to different mammography datasets and other medical imaging modalities. Additionally, incorporating longitudinal patient data, such as changes in breast tissue composition and tumor progression over time, could further enhance the clinical utility of image synthesis and its application in diagnostic tools and medical training. Also, the generated images can be employed in patient education, enabling healthcare professionals to visually explain various breast cancer types and stages to patients. This may lead to better understanding and compliance with treatment plans, ultimately improving patient satisfaction and outcomes.

In conclusion, the application of diffusion models in generating medical imaging data has the potential to significantly impact the field of medical image research by expanding training datasets and facilitating large-scale investigations. This novel approach offers the opportunity to create diverse and clinically relevant images without compromising patient privacy, thus enabling researchers to develop more accurate diagnostic tools, enhance medical education, and promote the advancement of innovative imaging technologies. The ongoing exploration and refinement of diffusion models will be crucial in shaping the future of medical imaging research, ensuring both improved patient outcomes and the ethical use of imaging data.

## METHODS

### Training the text-to-Image diffusion models using Dreambooth

Starting with multiple distinct sets of medical images we fine-tuned a text-to-image diffusion model that was trained using input images associated with a textual cue, which includes a distinct textual identifier for the class type (e.g. “An [xyz] mammogram”).

The models were trained using DreamBooth implementation of stable diffusion using distinct sets of medical imaging data. The framework is implemented using Python 3 and relies on the PyTorch library for deep learning tasks. The DreamBooth model consists of a denoising score matching (DSM) architecture with normalization layers, a U-Net based generator, and several residual blocks. The model is designed to perform image synthesis through a diffusion process. The architecture is configured to ensure stable and efficient training.

The initial model weights were obtained from the pretrained model “runwayml/stable-diffusion-v1-5” from Huggingface (https://huggingface.co/)^38^, which utilizes the diffusers library (https://github.com/huggingface/diffusers)^39^. The training configuration uses a seed value of 1337 and a resolution of 512×512 pixels for image generation. The training batch size is set to 1, and the model employs mixed precision training with 16-bit floating-point arithmetic (fp16) for improved computational efficiency. The Adam optimizer is used with 8-bit encoding, a learning rate of 1e-6, and a constant learning rate scheduler. No learning rate warmup is utilized. The model was trained on 50 class images per epoch, with a maximum of 800 training steps. Gradient accumulation was set to 1 step. The training process incorporates prior preservation with a prior loss weight of 1.0. During training, sample images are generated using a batch size of 4 and distinct prompts (e.g. “xyz mammography.” The concepts list for training is provided in a JSON file “concepts_list.json.” The model saves checkpoints at intervals of 10,000 steps.

The training process involves a list of concepts, which can be customized by adjusting the --max_train_steps parameter. In this example, the concept list consists of a single concept related to brain MRI images. An instance prompt is “brain MRI image,” and the class prompt is “brain MRI.” Each concept’s instance and class data directories was created using the os.makedirs function. The concept list was saved in a JSON file “concepts_list.json” for use during training. This method allows for the exploration of multiple concepts and their impact on the model’s performance, providing insights into the diffusion process and image synthesis.

### Diffusion Process

The diffusion process is governed by a stochastic differential equation (SDE) that involves a continuous-time Markov process. The SDE is solved using the Euler-Maruyama method with discretization steps (timesteps) ranging from 0 to T-1. The study explores different noise schedules to understand their impact on the diffusion process.

### Inference and image synthesis

The DreamBooth model was used to generate images based on specified prompts. During the image synthesis process, the diffusion process is reversed by iteratively applying the learned denoising function to a sequence of noise-corrupted images. The resulting image is obtained at the end of the reverse process. This procedure is performed with varying noise schedules to analyze the impact on image quality.

A random seed is set to ensure reproducibility of the results, with the seed value in this example being 52362. The main image generation parameters include the positive prompt, which is set to the specific model (e.g. “brain MRI image,”) and the negative prompt, which remains empty in this case. The number of samples generated is set to 4, while the guidance scale, which controls the strength of the guidance, is set to 7.5.

The image generation process was performed with a height and width of 512×512 pixels, using 24 inference steps for each image. The images were generated using a PyTorch pipeline with automatic mixed precision (“cuda”) and inference mode to optimize computational efficiency. The pipe() function receives the specified prompts, dimensions, and other parameters and returns the generated images.

Upon completion of the image generation process, the synthesized images are displayed or saved. This method allows for the exploration of the model’s performance in generating images based on given prompts and the impact of varying parameters, such as the guidance scale and number of inference steps. The reproducibility provided by the random seed ensures consistent results, facilitating the evaluation and comparison of the model’s performance under different conditions.

## DECLARATIONS

## ETHICS APPROVAL AND CONSENT TO PARTICIPATE

Not applicable

## CONSENT FOR PUBLICATION

All authors have read and approved the final version of this manuscript.

## Data Availability

Publicly available medical imaging data analyzed in this study:

<u>Brain MRI images</u>

Kaggle Brain Tumor Image Dataset^40,41^

https://www.kaggle.com/datasets/denizkavi1/brain-tumor

https://www.kaggle.com/datasets/mateuszbuda/lgg-mri-segmentation

https://wiki.cancerimagingarchive.net/pages/viewpage.action?pageId=5309188

<u>Lower-grade glioma (LGG) images</u>

lower-grade glioma (LGG) Segmentation Dataset^48^.

https://www.kaggle.com/datasets/denizkavi1/brain-tumor

<u>Mammography images</u>

CDD-CESM repository^49^

https://wiki.cancerimagingarchive.net/pages/viewpage.action?pageId=109379611

<u>NIH Chest X-rays</u>

ChestX-ray8 database^56^

https://www.kaggle.com/datasets/nih-chest-xrays/data

## AUTHORS’ CONTRIBUTIONS

B.L.K. conceived of the study, designed and carried out the experiments, analyzed the data, and drafted the manuscript.

## ACKNOWLEDGEMENTS

This work utilized the Wayne State University High Performance Computing Grid for computational resources (https://www.grid.wayne.edu/). This work was supported by Wayne State University and Karmanos Cancer Institute.

